# Comprehensive Hierarchical Classification of Transposable Elements based on Deep Learning

**DOI:** 10.1101/2024.01.27.577599

**Authors:** Yang Qi, Yiqi Chen, Yingfu Wu, Yanyan Li, Meihong Gao, Fuhao Zhang, Xingyu Liao, Xuequn Shang

## Abstract

Transposable elements (TEs) are DNA sequences capable of translocating within a genome. They constitute a substantial portion of eukaryotic genomes and play significant roles in genome evolution and gene regulation. The correct classification of these repetitive elements is essential to investigate their potential impact on genomes. Despite the existence of several tools for TE classification, they often neglect the importance of simultaneously utilizing global and local information for TE-type identification, resulting in suboptimal performance. Furthermore, these tools are not user-friendly due to the complex installation processes and numerous dependencies. In this study, we introduced a novel framework, CREATE, which leverages the strengths of **C**onvolutional and **R**ecurrent Neural N**E**tworks, combined with **A**ttention mechanisms, for efficient **TE** classification. Given the tree-like structure of TE groups, we separately trained nine models within the class hierarchy. Benchmarking experiments showed that CREATE significantly outperformed other TE classification tools. The source code and demo data for CREATE are available at https://github.com/yangqi-cs/CREATE. To facilitate TE annotation for researchers, we have developed a web platform, named WebDLTE, based on the CREATE framework. This platform employs GPU-accelerated pre-trained deep learning models for real-time TE classification and offers the most comprehensive collection of TEs for download. The web interface can be accessed at https://www.webdlte.nwpu.edu.cn.

## 1. Introduction

Transposable elements (TEs), also known as transposons or “jumping genes”, are DNA sequences with the capability to move or transpose within a genome [1]. TEs are ubiquitous across all eukaryotic kingdoms and make up a considerable fraction of their genomes. For instance, TEs make up approximately 45% [2] of the human genome and represent about 63% of the maize genome [3, 4]. Numerous studies have underscored the profound influence of TEs on genome evolution, genetic instability, and gene expression regulation [5-7]. With the widespread availability of whole-genome sequencing data from diverse species, unprecedented opportunities have been provided to investigate TE content and their relationships with genes. Owing to the presence of repetitive characteristics, TEs pose serious difficulties during the process of sequencing, genome assembly, as well as structural and functional annotation of genes [8]. However, TE discovery and annotation remain tricky tasks due to the polymorphisms of structures and variable lengths among TE sequences.

Numerous approaches have been developed for the identification of TEs in genomes, which can be generally divided into three primary categories: homology-based, signature-based, and *de novo* methods [9, 10]. In the homology-based manner, a BLAST-like tool is used as a search engine against the reference library. Censor [11] and RepeatMasker (http://www.repeatmasker.org/) are well-known approaches to homology-based TE annotation methods. Over the years, RepeatMasker has become a gold standard for TE annotation [12]. Nevertheless, these approaches thoroughly rely on the similarity between query sequences and the given annotated libraries, and they can only detect TEs that had been described previously. Signature-based approaches search for certain structural features, such as long terminal repeat (LTR), target site duplication (TSD), or primer-binding site (PBS), which can characterize specific TEs. For example, LTR_FINDER [13] and LTR_retriever [14] are designed based on this principle and aim to find LTR retrotransposons, while MITE-Hunter [15] and MITEFinderII [16] are tailored for the detection of MITEs. In this scenario, substantial and sufficient prior knowledge is required to identify a particular TE group. Moreover, signature-based methods are incapable of identifying TE types that exhibit a deficiency in primary structural features [17]. Since *de novo* approaches do not rely on prior knowledge of the structure or similarity to known TEs, and instead leverage the repetitive properties of sequences to identify TEs, they exhibit greater versatility compared to the two aforementioned algorithmic groups. Numerous programs have been developed based on the *de novo* concept, such as RECON [18], RepeatScout [19], RepARK [20], RepLong [21], and LongRepMarker [22]. It is worth noting that these approaches enable the recognition of any repetitive elements present in the genome to construct consensus sequences. In this fashion, a sophisticated classification tool is required to annotate specific types for these candidate TEs. Accurate classification of transposons is an essential step in genome annotation and offers an opportunity to unravel their roles in germline and somatic evolution [23].

TE classification is the process of categorizing TEs based on their structure characteristics and transposition patterns. The first TE classification system was proposed by Finnegan in 1989 [24]. In this classification proposition, TEs were divided into two major categories according to their transposition intermediate: class I elements (retrotransposons) employ RNA as an intermediate for transposition through a copy-and-paste mechanism, while class II elements (DNA transposons) mobilize via a cut-and-paste fashion. Since the discovery of a novel type of TEs that exhibits copy-and-paste mechanisms devoid of RNA intermediates, the prevailing two-class system has faced considerable scrutiny and questioning. In 2007, Wicker *et al*. proposed a unified hierarchical classification system that retained two main classes and distinguished TEs by the presence or absence of RNA intermediates [25]. In this taxonomy, class I TEs are further divided into five orders: the LTR order and four non-LTR retroelement orders, including DIRS, PLEs, LINEs, and SINEs. Meanwhile, class II transposons are separated into two subclasses, which reflect the number of DNA strands that are cut during transposition. Subclass 1 contains the elements that transpose through the pre-established cut-and-paste mechanism, whereas subclass 2 consists of Helitron and Maverick elements, which move by cutting only one strand of the donor.

Automated TE classification is a pivotal step in deciphering the essence, significance, and influence of TEs on genome organization and functionality. Homology-based and structure-based tools for recognizing TEs can be utilized to a certain extent for classification purposes, and there are also specialized classification tools available. REPCLASS [26] comprises three modules: the HOM module conducts the homology search in the reference library for each entry; the STR module employs various subroutines to detect structural characteristics like LTR, TIR, etc.; and the TSD module determines the potential target site duplications that flanking the insertion site of individual TEs. Eventually, the final step endeavors to compare and combine the outcomes of three modules, leading to a preliminary classification for each input sequence. PASTEC [27] not only investigates structural features and sequence similarities but also incorporates Hidden Markov Model profiles of conserved domains of proteins to infer types of TEs. Both of these methods utilize similarity-based and structure-based strategies, which rely on the completeness of the prior information database and involve an intricate methodological pipeline. Consequently, there is a pressing need to develop general tools that can accomplish TE classification more efficiently.

In the past few decades, machine learning technologies have been widely applied in various research fields. TEclass [28] is the first learning-based tool for TE classification, which leverages support vector machines (SVMs) for categorization based on oligomer frequencies. As a specialized branch of machine learning, deep learning excels in extracting complicated patterns from the raw data. Significantly, deep learning-based models have consistently surpassed established machine learning techniques in numerous scientific fields. DeepTE [29] and TERL [30] are two current state-of-the-art deep learning-based approaches for TE classification. DeepTE employs CNN models and trains eight models to classify unknown sequences into superfamilies and orders. Both DeepTE and TEclass utilize *k-mer* counts as input features. Despite the *k-mer* frequency has been successfully applied to many approaches, it could still lose some original information of the initial sequences. In TERL, the authors proposed a novel solution to transform one-dimensional sequence data into two-dimensional space data and applied it to the CNN model. However, TERL overlooks the hierarchical categorization scheme of TE groups. ClassifyTE [31] is a specified hierarchical classification approach, which constructs an elaborate hierarchical dataset and develops a stacking-based machine learning framework for TE classification. However, the rationale for annotating elements as non-leaf nodes is not yet clear. It remains uncertain whether this is due to the characteristics of the sequence itself or the result of technical errors. Recently, we noticed a preprint manuscript called TEclass2 [32], which utilizes the Longformer model for TE classification. This method proposes an all-in-one classifier that assigns sequences into 16 superfamily categories, but it lacks many valuable TE classes, like DIRS and PLE. It ignored the hierarchical scheme of TE class and restricts sequences classified into specific groups, which is inconsistent with the actual evolutionary relationships of transposons. In addition, the Longformer architecture used in TEclass2, while capable of handling large-scale features, has high hardware (e.g., GPU) requirements, making it unsuitable for ubiquitous computing environments.

Current learning-based algorithms for TE classification predominantly concentrate on the global feature representation of sequences. They often overlook the crucial structural information derived from both ends of TEs. This oversight limits the sensitivity of these algorithms. In this study, we presented an innovative approach, CREATE, which leverages the advantages of combining **C**onvolutional and **R**ecurrent Neural N**E**tworks (CNN and RNN) along with **A**ttention mechanisms to achieve efficient **TE** classification. In CREATE, nine models were meticulously trained according to the class hierarchy (Figure 1A). For each classifier, the CNN module was utilized to acquire information from the global *k-mer* frequency, while the RNN module was employed to extract the local potential homologous information through the manually crafted terminal sequences. In addition, the attention mechanism was used to integrate the output features of the CNN and RNN modules (Figures 1B and C). Therefore, both the global and local characteristics of transposons can be depicted in the classification process. Due to the nature of TE datasets being generated, the classification issue can be assumed as a non-mandatory leaf node prediction problem [31]. This strategy allows instances to be classified to any level and achieves an end-to-end criterion for hierarchical TE classification in the testing phase.

**Figure 1.**
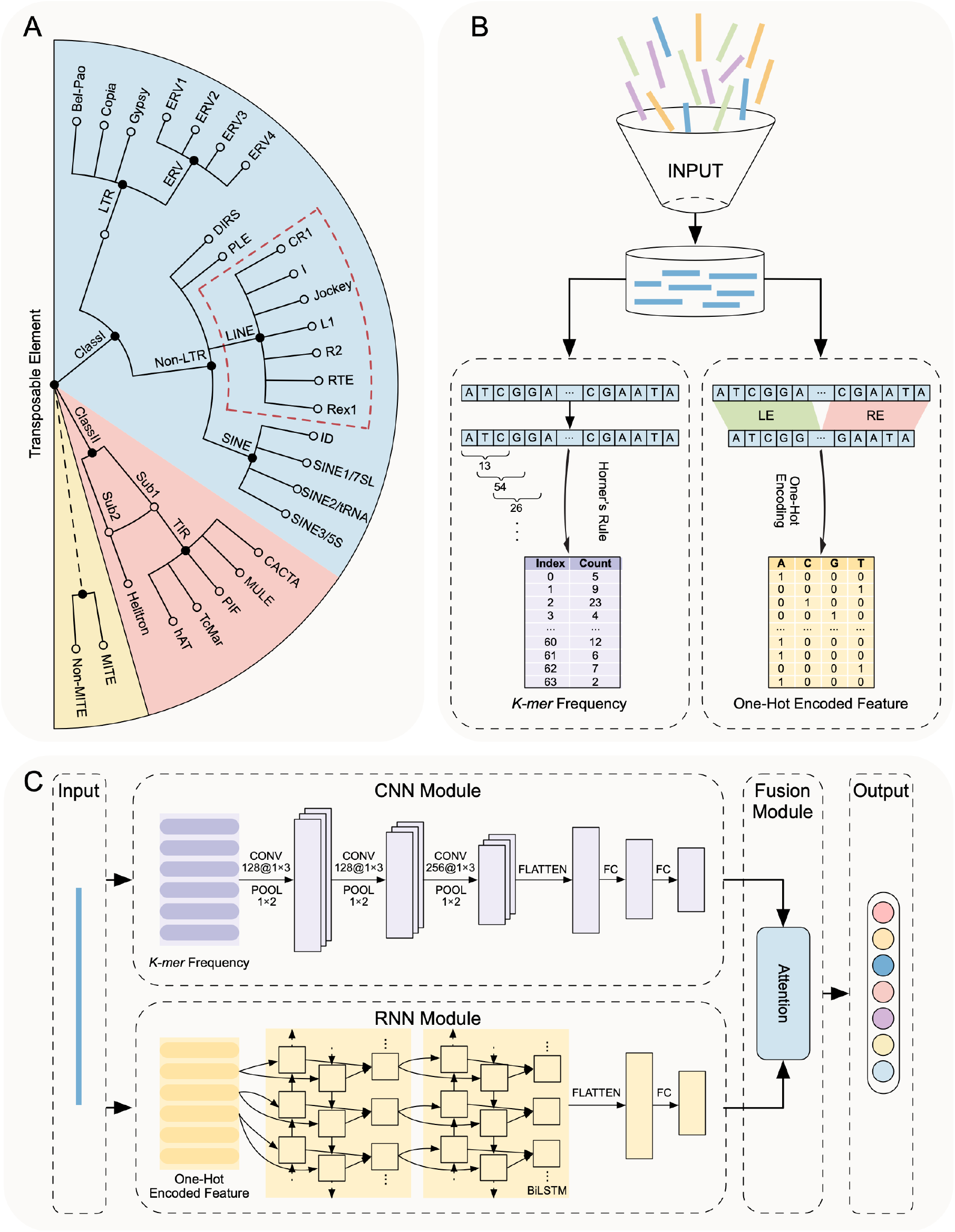
The schematic diagram of CREATE. (A) The relationship between classification models and TE groups. In CREATE, nine parent nodes (solid circles) in the circle tree were trained with multi-class classifiers separately. (B) In the sequence feature extraction phase, Horner’s rule is utilized for efficient *k-mer* frequency extraction, while double terminal sequences are represented through one-hot encoding. LE: left end. RE: right end. (C) The framework of the attention-based hybrid CNN-RNN architecture for each classifier. CONV: convolutional layer. POOL: max pooling layer. FC: fully connected layer. BiLSTM: bidirectional long short-term memory.

To streamline the process of TE annotation for researchers, we have designed a web interface, WebDLTE. This platform provides the advantage of performing high-precision online classification of TE sequences, leveraging pre-trained models developed by the CREATE method. Furthermore, WebDLTE functions as a comprehensive database for regenerated TE sequences, enabling users to seamlessly access and download their desired sequences. The interface facilitates unrestricted search capabilities, allowing users to search by either species name or transposon type. Additionally, the platform also provides the function of analyzing the evolutionary relationships of transposons, which is helpful for studying biological evolution.

### Highlights

Compared with the prevailing TE classification methods, this study presents several notable advantages, which can be outlined as follows:

i. **Provide a new perspective on sequence feature representation and fusion**. The CREATE method considers two key factors when determining TE types: the global content of oligomers and the local bipartite structure of TE sequences. Furthermore, it employs an attention mechanism that enables the model to adaptively learn the importance of different features for various TE types, thereby facilitating a more efficient integration of these features.
ii. **Provide efficient feature extraction and high-precision TE hierarchical classification**. The traditional approach of sequential traversal for calculating *k-mer* counts suffers from inefficiencies in time complexity. However, Horner’s rule provides a solution by enabling the conversion of the *k-mer* sequence into a decimal number from its quaternary representation, significantly accelerating our feature extraction process in CREATE. Moreover, CREATE offers users a high-precision, hierarchical categorization of TEs, with a user-defined threshold in the testing phase, ensuring that category labels are logically coherent and not obligatory. Comprehensive experimental results validate that the CREATE framework surpasses existing cutting-edge methods in TE classification.
iii. **WebDLTE, an important extension of the CREATE method, stands as the pioneering website to employ deep learning for real-time TE classification**. WebDLTE, the inaugural website to utilize GPU-accelerated pre-trained deep learning models for online TE classification, serves as an innovative platform that greatly simplifies TE annotation for researchers, thus paving the stage for novel investigations into the biological functions of TEs.
iv. **A broader range of TE consensus sequences is provided in WebDLTE**. While existing TE databases like Dfam and RepBase are typically kingdom specific, WebDLTE offers a comprehensive collection of TE consensus sequences from a diverse range of 315 eukaryotic species, including animals, plants, fungi, and more. With a total exceeding 932,944 sequences, it currently boasts the most complete collection of TEs.

## 2. Methods

### 2.1 Overview

Considering the tree-like structure of TE groups, a novel deep learning-based method, CREATE, was proposed for the hierarchical classification of TEs. In CREATE, the local classifier strategy was adopted [33], and nine attention-based hybrid CNN-RNN architectures were trained for each parent node in the class hierarchy. Specifically, for each model, *k-mer* counts were employed as global characteristic input to the CNN module, while the handcrafted double terminal sequences were extracted as local feature input to the RNN module. Then an fusion module based on attention techniques was utilized to weigh the features extracted by CNN and RNN. For unknown TEs, CREATE can assign them to any class level in the hierarchical system through a non-mandatory leaf node prediction strategy.

A detailed description of each portion is provided in this section. **Section 2.2** describes the feature extraction approach. The details of each module in the framework, including CNN, RNN, and attention mechanisms, are outlined in **section 2.3**. Subsequently, the hierarchical classification strategy in the testing phase is introduced in **section 2.4**. The last subsection presents the evaluation metrics employed for performance comparison.

### 2.2 Sequence feature extraction

The success of machine learning algorithms hinges greatly on the quality of feature representation. In the field of bioinformatics, *k-mers* have gained significant traction due to their ability to depict the signature of relative distribution underlying the nucleotide sequences. In the CREATE framework, the length of *k-mer* frequency is *l* = 4^*k*^ where *k* is the size of *k-mers*. To overcome the time complexity introduced by the conventional method of sequential traversal, Horner’s rule was adopted as a time-efficient way to compute *k-mer* frequencies [34, 35]. Specifically, different nucleotides A, C, G and T in the *k-mer* are regarded as distinct digits 0, 1, 2 and 3. Each sequence is linearly scanned by a window of *k* letters, where the letters within each window are transformed into numerical values. These values are then converted from a quaternary representation to decimal numbers, which serve as indices in the hash table (Figure 1B). This strategy significantly accelerates the feature extraction process. In CREATE, the *k-mer* counts were preprocessed as the input layer of the CNN module. The performance of different *k-mer* sizes was analyzed (Supplementary Table S1) and the optimum value was selected as the final parameter for the downstream classification tasks.

Despite traditional *k-mers* are useful in feature extraction, they fail to capture intrinsic patterns and motifs in the TEs [36]. Inspired by the fact that double terminals of transposons hide abundant information about TE types, such as LTR and TIR, we extracted important sequence features from Both Ends (BE) of the TEs. Specifically, the sequences were truncated from the Left End (LE) and the Right End (RE) with particular length. Then the handcrafted LE and RE sequences were concatenated as a whole sequence (Figure 1B). Since RNNs are well-suited for processing sequential data, an RNN module was employed to extract hidden features from sequence encodings. The results of different lengths were tested (Supplementary Table S2) and the proper number was selected to train the model.

### 2.3 Attention-based CNN-RNN architecture

#### 2.3.1 The CNN module

CNN is one of the most widely used deep learning architecture. It typically includes three types of hidden layers: convolutional layers, pooling layers and fully connected layers. The convolutional layer is the core block of CNN, and it utilizes several kernels to execute convolution operations on the input data. Mathematically, the convolution operation can be expressed as follows [37]:

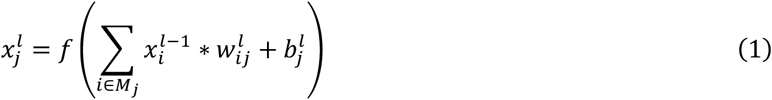

where 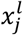 denotes the channel *j* in the current layer *l, M*_*j*_ represents a selection of the input channel, 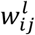 is the weight matrix of convolution, 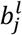 is the bias vector, and *f*(·) represents the activation function. Then the pooling layer is applied after the convolutional layer. The function of the pooling operation is to perform subsampling and prevent overfitting by reducing the size of features. The pooling operation can be represented as follows:

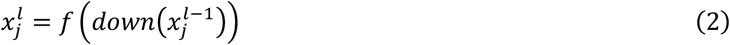

where *down*(·) refers to a subsampling function. Max pooling is one of the commonly used pooling techniques, and this strategy was utilized in our CNN module. Finally, fully connected layers are employed to map the extracted features into the final output and the output of this layer is defined as:

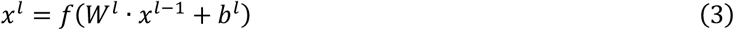

where *W*^*l*^ is the weight matrix and *b*^*l*^ is the bias.

In the CNN module of the CREATE framework, there are three convolutional layers, three max pooling layers and two fully connected layers (Figure 1C). ReLU was adopted as the activation function in all hidden layers, and a dropout value of 0.5 was set after the last convolutional layer and the two fully connected layers. In the training process, categorical crossentropy was used as the loss function, and the ADAM algorithm was adopted as the gradient-based optimizer with a learning rate of 0.0005.

#### 2.3.2 The RNN module

The RNN model has been widely used to recognize the sequential characteristics in data. However, RNN has limitations in long-term dependencies and can lead to vanishing gradient problem. Long short-term memory (LSTM) is a special type of RNN that incorporates three types of additional gates, including input gate *i*_*t*_, forget gate *f*_*t*_, output gate *o*_*t*_, as well as a cell state *C*_*t*_ to memorize a longer sequence of input data. LSTM usually leads to better performance compared with the basic RNN [38]. At a time *t*, the components within the LSTM are updated as follows [39]:

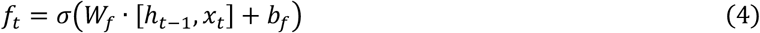

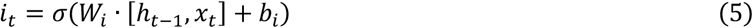

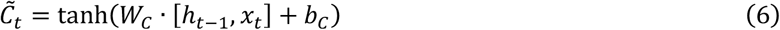

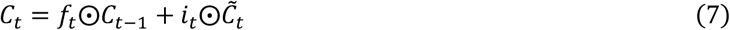

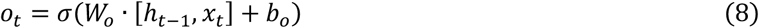

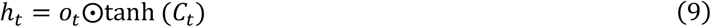

where *σ*(·) and tanh (·) are sigmoid and tanh activation functions respectively. *W*_*f*_, *W*_*i*_, *W*_*c*_ and *W*_*o*_ are the weight matrices of the gates or the cell. *b*_*f*_, *b*_*i*_, *b*_*c*_ and *b*_*o*_ are bias. ⨀ represents the elementwise multiplication and *h*_*t*_ stands for the hidden state.

Additionally, the bidirectional LSTM (BiLSTM) is an extension of the standard LSTM, and it enables additional training by traversing the input data twice. The hidden state *h*_*t*_ of BiLSTM contains the output of forward LSTM 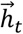 and backward LSTM 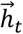:

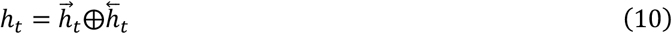

where ⨁ denotes the summation function to sum the forward and backward output components. In general, BiLSTM extends the prediction capability of LSTM [40].

Here, we employed two stacked BiLSTM layers in the RNN module and set 128 and 64 units for each LSTM, respectively. Then the BiLSTM layer was flattened and fully connected to a layer with 64 hidden units (Figure 1C). We also used the categorical crossentropy as the loss function in the CNN module, and the ADAM approach was employed as the optimizer with a learning rate of 0.0005.

#### 2.3.3 The Fusion module

In the framework proposed in this study, the CNN and RNN modules were trained through a parallel mode. Thus, both global and local information could be extracted by these two networks. Then, we leveraged the attention mechanism and effectively combined the output layers of CNN and RNN. In this manner, it is possible to dynamically allocate appropriate weight for these two modules [40]. We assumed the feature obtained by CNN or RNN as *h*_*s*_ similar to features at each timestamp *s* in the typical attention model, with *W* representing the weight matrix. *h*_*s*_ was then followed by a fully connected layer to obtain the hidden representation *e*_*s*_. Finally, a softmax function was employed to obtain the normalized importance weight, denoted as *α*_*s*_. Subsequently, the context vector *c* was computed using the weights *α*. The above procedure can be defined as:

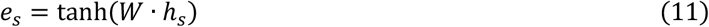

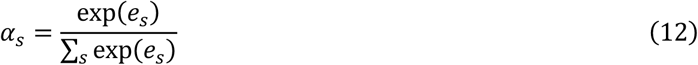

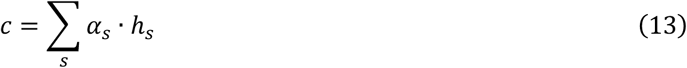

Through the attention mechanism, CREATE can dynamically allocate suitable weight for CNN and RNN modules and allocate attention to the appropriate component.

### 2.4 Hierarchical classification

TE-related hierarchical categorization schemes can be considered as a hierarchical classification problem. Contrasting with other hierarchical methodologies, the strategy of the local classifier per parent node involves training a multi-class classifier for each parent node within the class hierarchy. This is done to differentiate its child nodes, thereby mitigating problem complexity by fragmenting the original issue into smaller subproblems [42]. In this manner, nine classification models are trained in the CREATE framework (Figure 1A).

During the testing phase, this approach can also be coupled with a top-down prediction approach. Due to the characteristics of TE datasets being generated, TE classification can be considered a non-mandatory leaf node prediction problem, which allows instances to be classified to any level of the hierarchy. Here, we utilized a probability threshold as the stopping criterion to determine whether an example belongs to the current node or should be given to the current node’s child-node classifier. More precisely, if the confidence score of the classifier at the given class is lower than the threshold, the classification stops at the current node. Otherwise, the model the of sub-node with the highest probability will be employed as a new classifier (Figure 2). This greatly enhances the authenticity and reliability of the model’s inference for unknown sequence labels.

**Figure 2.**
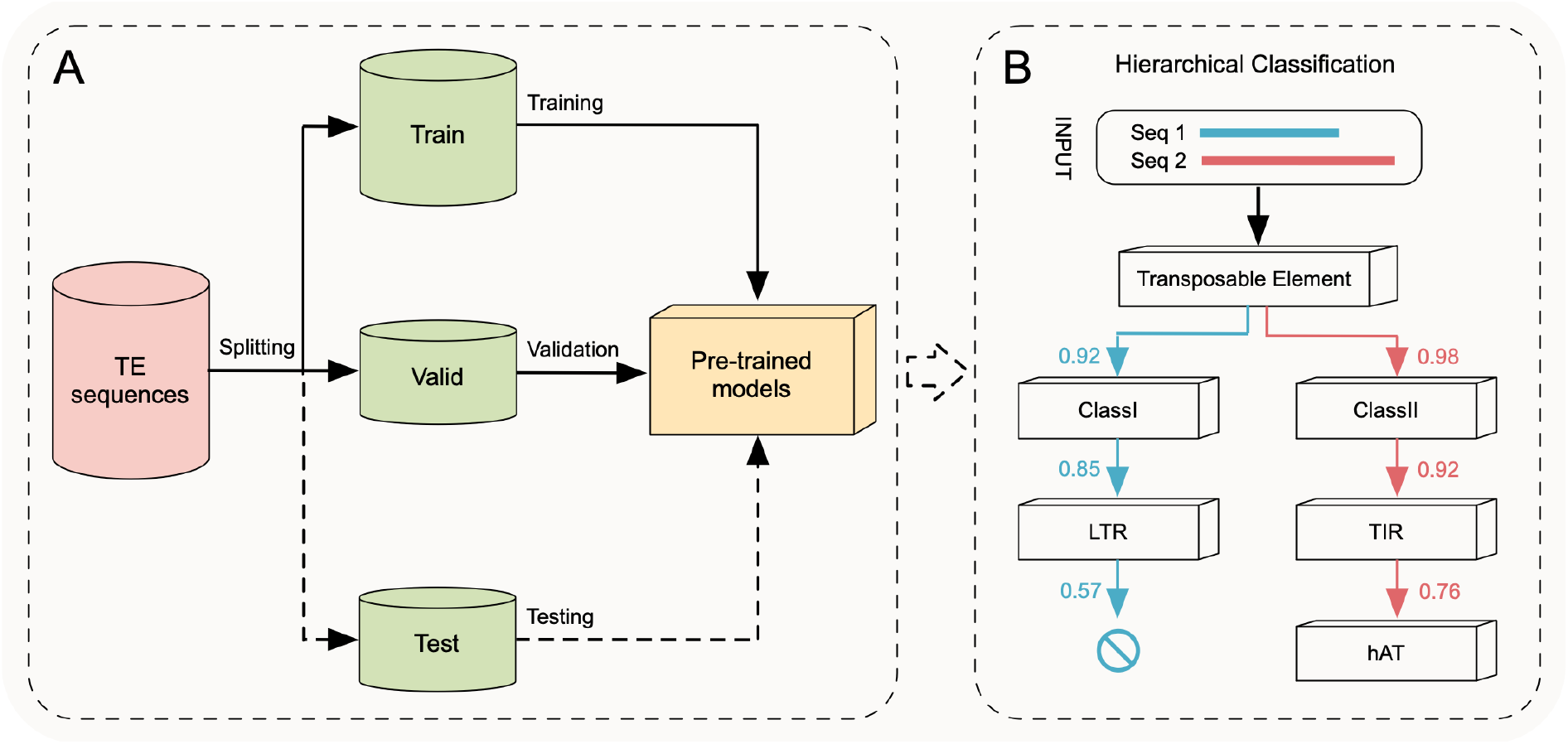
Depiction of the top-down approach for class prediction. (A) The preprocessed TE sequences are split into three sets (train, validation, and test). The testing portion of the model is indicated by dotted lines. (B) Illustrated two template sequences in the hierarchical classification pipeline, with 0.6 as the probability threshold. Seq 1 stopped at the non-leaf node, LTR, under the probability 0.57 (less than 0.6), whereas was seq 2 classified as a leaf node hAT.

### 2.5 Evaluation metrics

To evaluate the classification performance of classifiers, we adopted multiple classification metrics, including accuracy, precision, recall, F1-score, and Matthews Correlation Coefficient (MCC). These metrics are specified as:

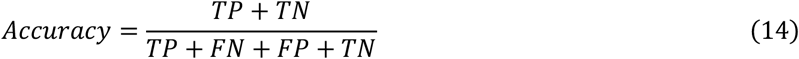

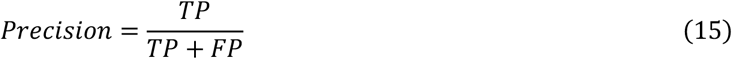

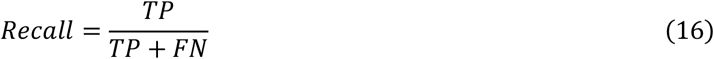

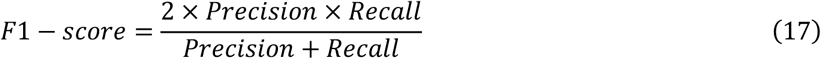

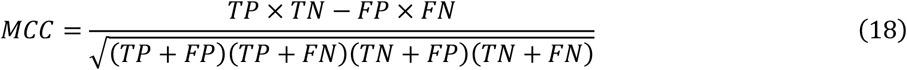

where TP (true positive) and TN (true negative) represent the number of correctly classified samples in the positive and negative classes, respectively. FP (false positive) and FN (false negative) stand for the number of incorrectly classified samples in the positive and negative categories, respectively. The ROC curve provides a graphical representation of the classifier’s performance, and the area under the ROC curve (AUC) provides an aggregate measure to compare the performance of different approaches. Considering the class imbalance problem in the original dataset, macro-average scores were calculated by averaging scores for each class, and this metric treats all classes equally to evaluate the overall performance of the classifier.

Providing high-precision end-to-end automatic hierarchical classification is a highlight of our proposed framework. To evaluate the performance of hierarchical approaches, the extended versions of standard metrics were calculated by estimating the classification results at each hierarchical level. Hierarchical precision (hP), hierarchical recall (hR), and hierarchical f-measure (hF) were used as the evaluation as follows [33]:

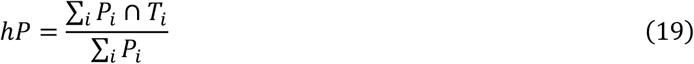

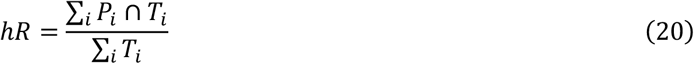

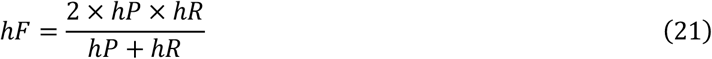

For a test sample *i, P*_*i*_ is the set consisting of the most specific predicted label and its ancestor labels, and *T*_*i*_ is a set composed of the most specific true label and its ancestor labels.

## 3. Results

### 3.1 Data collection and preprocessing

To construct a highly credible TE dataset, we collected sequences from 13 widely used repeat databases (Table 1). The non-TEs abnormal sequences were removed, such as unlabeled, satellites, and RNA sequences [43]. Then the sequences in these databases underwent standardized preprocessing as follows: (i) Filtering out sequences with a length less than 80 bp; (ii) The sequences with more than 20% non-canonical characters were also removed (e.g., non-A, C, G, or T); (iii) Only non-identical TE sequences were retained for subsequent studies. Due to the extremely large number of sequences in the APTEdb database, which is of a different magnitude compared to other databases, we randomly selected a specific number of sequences from our database for a particular TE category to avoid the excessive influence of this database on the generalization performance. This quantity of TEs is determined by taking the minimum value between the number of sequences in the specific category in the APTEdb database and half of the total number of sequences in the same category across all other databases. (iv) The consensus sequences generated in the previous step were formatted using the following naming convention: “ID|TE type|Species information”. To annotate Class II elements into MITE and non-MITE TEs, we manually curated the MITE and non-MITE datasets. TEs annotated as MITE in databases were labeled as “MITE”, and the remaining DNA sequences were considered as candidate non-MITEs. The authenticity of candidate non-MITEs were further assessed based on the sequence length and protein domains.

**Table 1.**
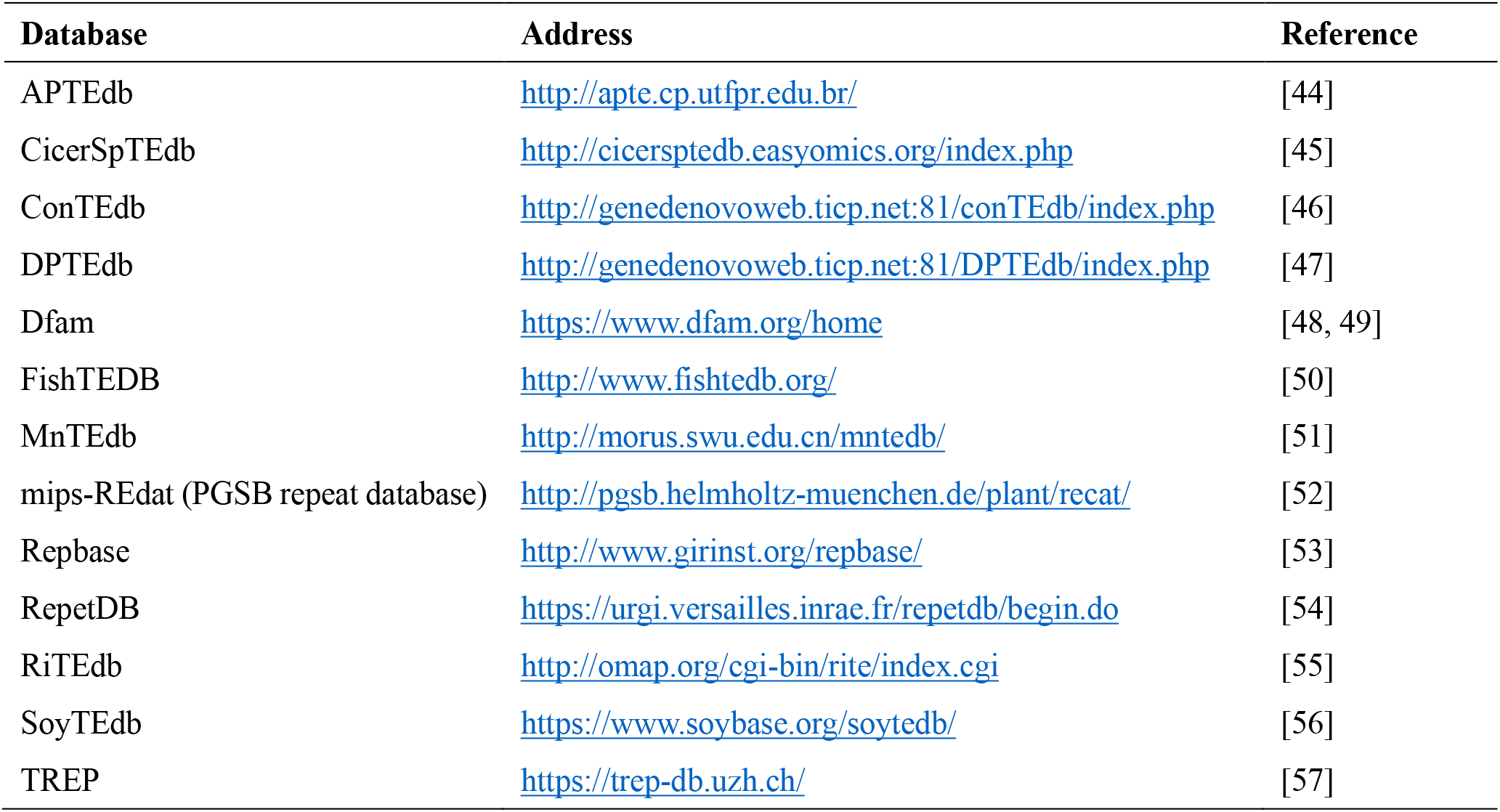
The repetitive sequences from 13 commonly used databases were collected and standardized. The preprocessed TEs serve as training data in the CREATE framework.

As far as we know, this dataset is the most extensive compilation of meticulously labeled TE sequences available to date. Encompassing a diverse array of TE orders and subfamilies, this comprehensive collection provides invaluable resources for studying the intricate dynamics and evolutionary significance of these genetic elements. To facilitate user accessibility, the TE consensus sequence was regenerated based on the similarity of sequences. Initially, the TE sequences belonging to species-specific superfamilies were clustered using CD-HIT [58], and then the consensus sequences were obtained by aligning the clustered TE sequences through abPOA [59]. By utilizing the comprehensive dataset at hand, we can delve deeper into the analysis of these sequences. This allows us to gain valuable insights into the diverse repertoire of TEs, thereby enhancing our understanding of the dynamic interplay between TEs and the forces that shape evolutionary trajectories ( see Figure 3). WebDLTE provides the regenerated consensus sequences that are labeled, standardized, and encompass a diverse range of species.

**Figure 3.**
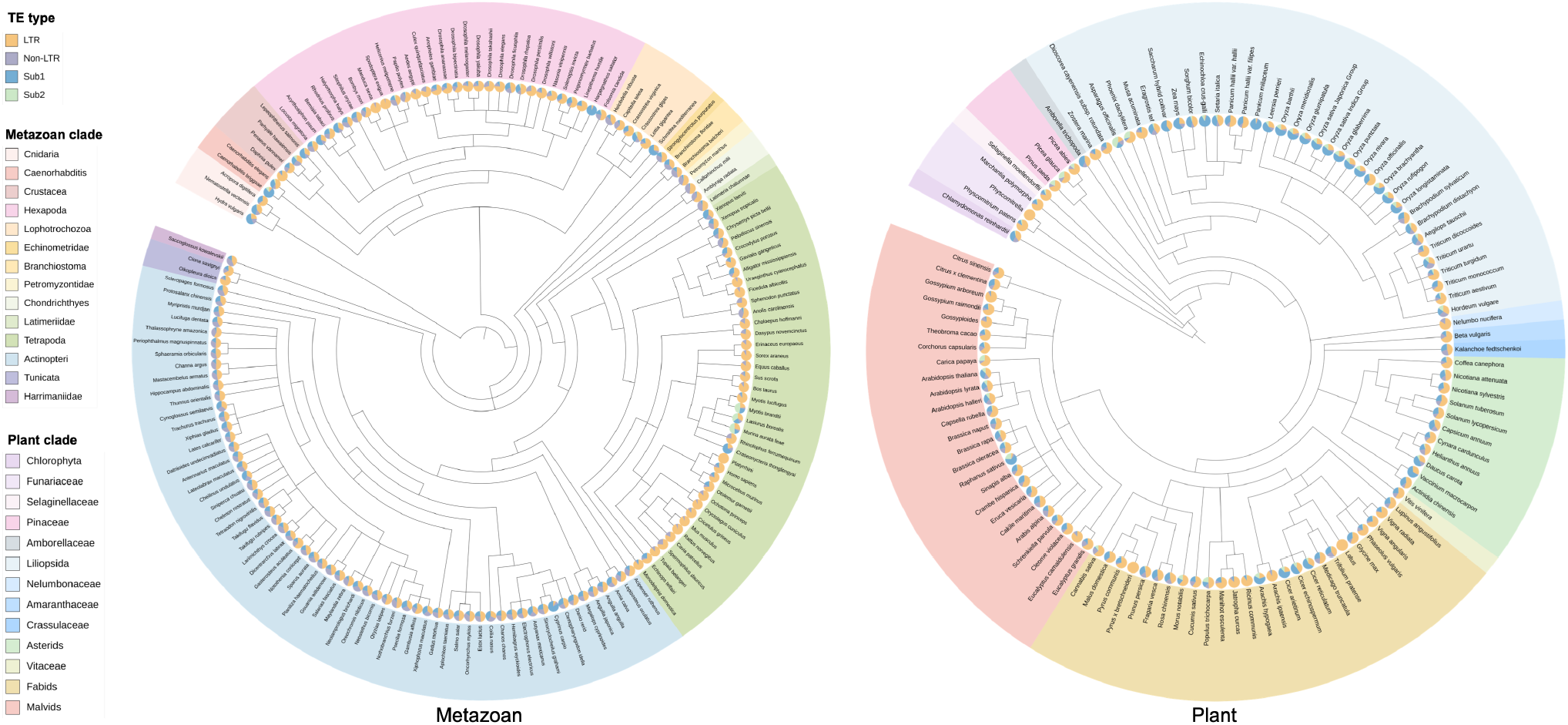
Phylogenetic trees of metazoans and plants constructed from TE sequences provided by WebDLTE. Pie charts in each branch correspond to profiles of four TE subclasses, including LTRs and non-LTRs in class I transposons, and subclasses 1 (Sub1) and 2 (Sub 2) in class II elements.

To validate the model’s generalization performance for transposons in unknown species, we utilized only TE sequences in metazoans and plants in the training phase. Additionally, 5,020 sequences of fungi were employed to test the model’s classification capability. Therefore, a total of 932,944 TEs were left for the downstream classification task (Supplementary Table S3). To estimate the prediction capability of different deep learning-based classifiers, we adopted a stratified sampling strategy and split the dataset into training (80%), validation (10%), and testing (10%) datasets (Table 2). TEs play significant roles in driving the adaptive evolution of plants [60]. As a model organism for plant biology and agricultural research, rice serves as an excellent choice among cereals for genomic studies owing to its compact genome size and diploid nature [61]. For repetitive sequences in the rice genome, the authors of EDTA manually curated a high-quality TE library as the benchmark dataset [62]. There are 2,431 sequences in the reference library to verify the performance of the hierarchical classification pipeline proposed in this study.

**Table 2.**
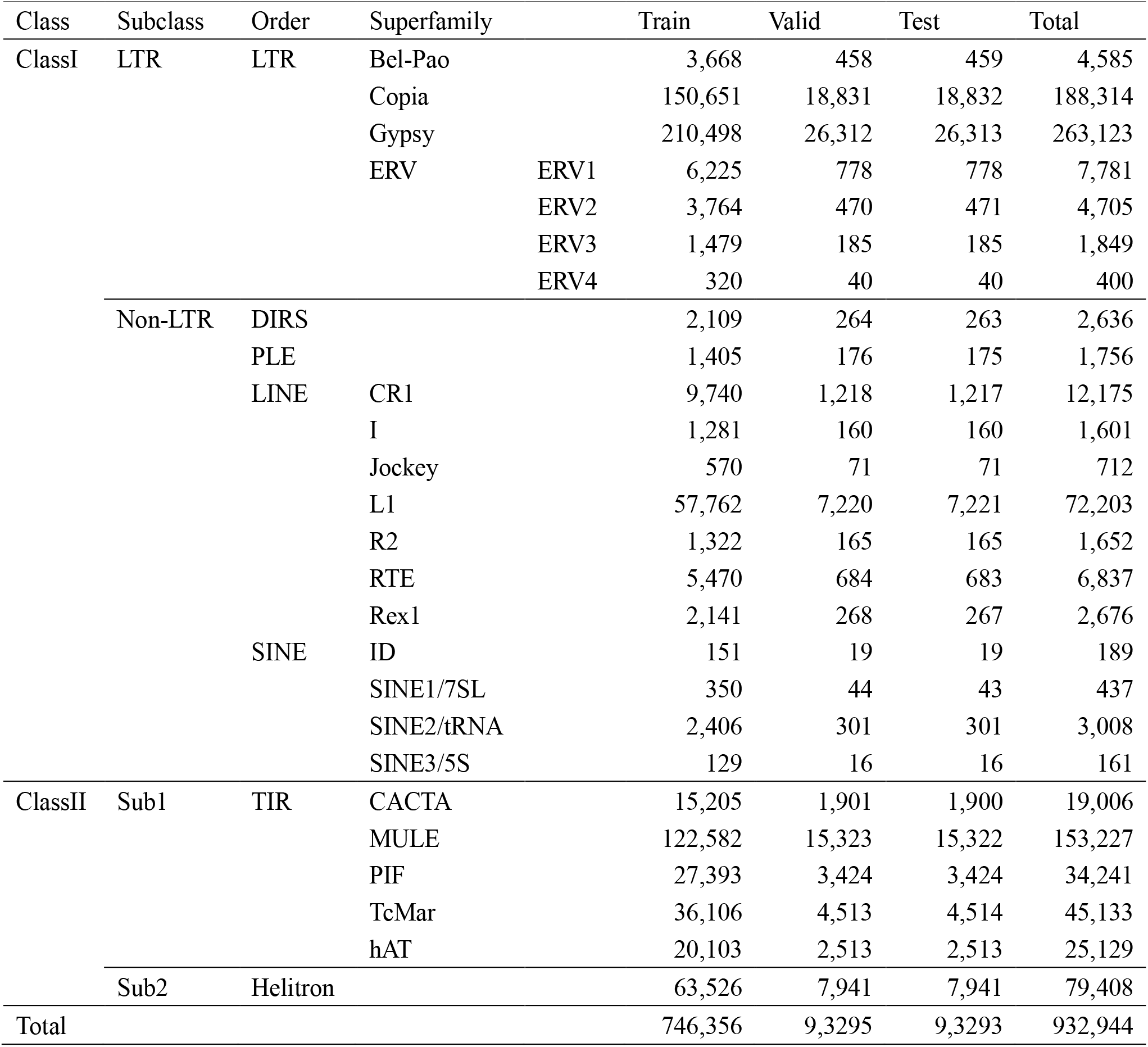
The hierarchical structure of TEs, along with the count of sequences allocated for training, validation, and testing datasets across various groups.

### 3.2 CREATE outperforms other TE classification tools

In the training process of CREATE, nine classifiers were trained separately on the basis of the class hierarchy. As mentioned above, Horner’s rule was employed as an efficient way for calculating *k-mer* frequency during the feature extraction phase. As shown in Figure 4A, with the increase in data volume, there is a remarkable reduction in the time required for feature extraction. For the model handling the largest dataset, specifically the Transposable Element model that includes all TE sequences, the traditional method took almost two hours for feature extraction. However, employing Horner’s rule expedited the process, completing it in merely one hour. This signifies that the employment of this method greatly enhances the operational efficiency of the model.

**Figure 4.**
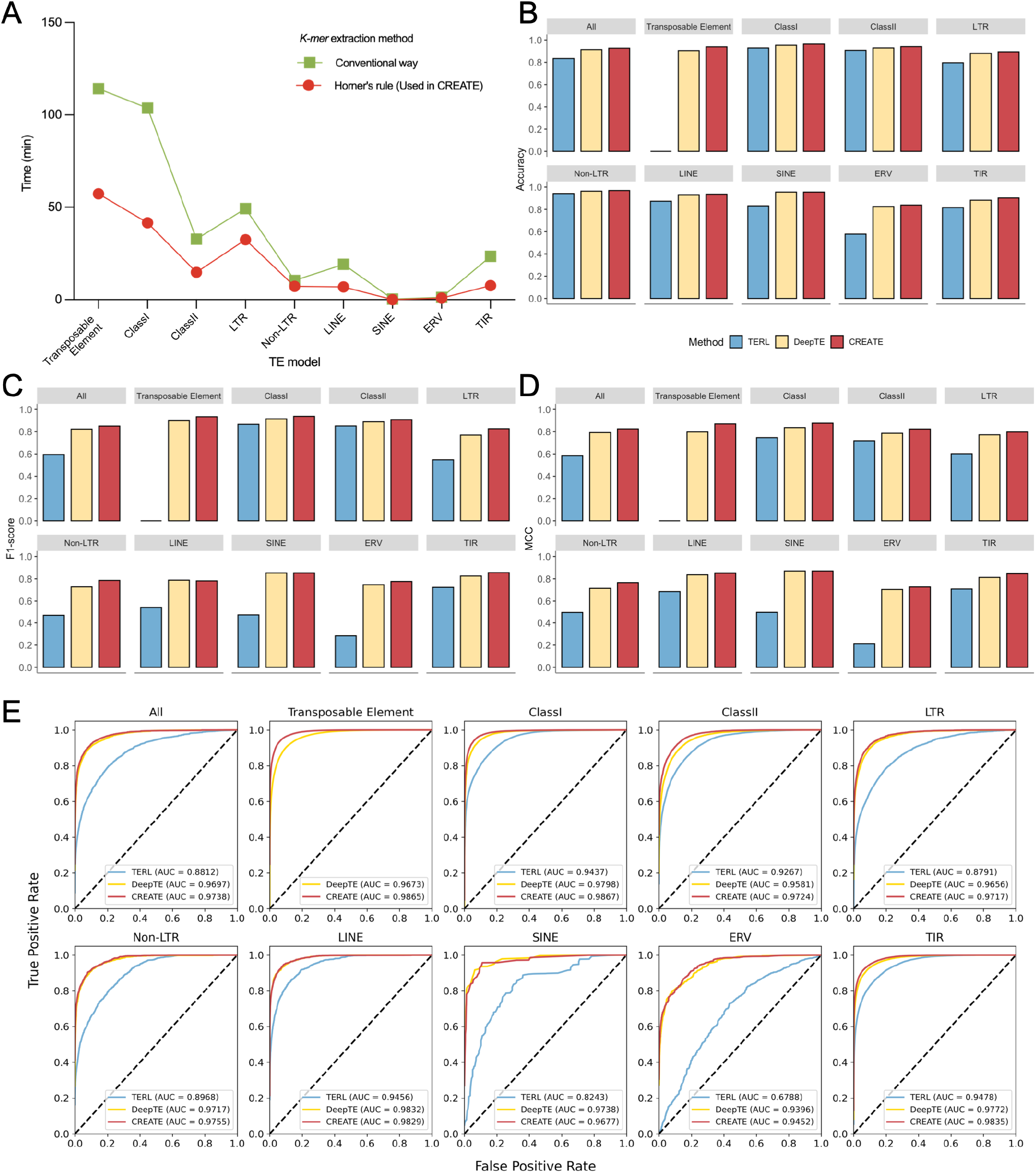
Performance comparison between different TE classification tools. (A) The runtime of different *k-mer* extraction methods, where Horner’s rule is adopted in the CREATE approach. (B-D) Performance of macro-average Accuracy, F1-score, and MCC for different tools. The bar plot in the All model represents the mean value across nine models for different methods. (E) ROC curves were obtained on the test dataset for different tools, where ROC curves in All models represents the macro-average plot across nine models for different methods.

To assess the performance of each model trained within the CREATE framework, we executed two additional deep learning-based TE classifiers, namely TERL and DeepTE, utilizing their default settings. Compared with other methods, CREATE achieved the best results in almost all model categories, only slightly inferior to DeepTE in a few exceptional groups of LINE (Supplementary Table S4). The macro-average scores of various metrics were also calculated for different classification tools (Figure 4B-D). CREATE obtained the best performance compared with other TE classification tools, and its values are higher than 0.850 in most models. The performance of these methods was further analyzed based on the average ROC curve. Figure 4E clearly illustrates that CREATE outperformed all other models, demonstrating its superior performance. This indicated that CREATE obtained a higher true positive rate for the same false positive rate compared with other methods.

To evaluate the classification performance of each component in CREATE, the separate experiments were also conducted to demonstrate the effectiveness of each module in CREATE. CREATE-CNN indicates the approach which leverages the CNN module of CREATE as the classification tool. Likewise, CREATE-RNN means applying the RNN module of CREATE as the classifier to classify TEs. We conducted experiments on high-quality subdatasets constructed TEs from Repbase and mips-REdat. The results demonstrate that, compared with other CNN-based classification approaches, CREATE-RNN exhibited vastly superior to classify TEs in imbalanced classes (Supplementary Table S5 and Figure S1). It is worth noting that CREATE-RNN produced better performance than CREATE-CNN in both Class II and TIR models, which compensate for the shortages of CNN classifier. In addition, CREATE-RNN greatly enhanced the performance of CREATE and outperformed other classification tools in most minority classes, which suggested that CREATE is more robust, and the class-imbalance problem has a relatively small effect on it. In addition, TERL performed poorly compared with other methods. These may be caused by the preprocessing step of TERL, which robustly padded all sequences into the same length as the longest one and introduce a lot of noise information.

For different classification tools, the non-mandatory leaf node prediction strategy was conducted as proposed in section 2.4. Notably, ClassifyTE is a specialized hierarchical classification approach that constructed the hierarchical dataset and added another subclass for each parent node to represent itself. Since the elements collected in our experiment are all TE sequences with specific information about the superfamily (leaf node), which is inconsistent with the fundamental assumptions of ClassifyTE, we simply employed the trained models provided in ClassifyTE,. For other methods (TERL, DeepTE, CREATE-CNN, CREATE-RNN, and CREATE), the final models were trained by all transposons from the combined Repbase and mips-REdat datasets. We calculated the hP, hR, and hF scores for each method using the probability threshold range from 0.50 to 0.95 with 0.05 as the step size. CREATE had an overall better performance than other methods in both hP, hR, and hF metrics (Supplementary Figure S2). In addition, CREATE is a parameter-insensitive approach, implying that it exhibits stronger robustness in TE classification tasks.

### 3.3 Web interface of WebDLTE

To facilitate TE annotation for researchers, a website, WebDLTE, is designed to offer online TE classification based on the pre-trained models produced by CREATE method. To the best of our knowledge, this is the first website that employs GPU-accelerated pre-trained deep learning models in the backend to offer online TE classification. This innovative platform facilitates sequence annotation for researchers, thereby unlocking new avenues for exploring phylogenetic relationships. WebDLTE consists of three main functional pages. The SEARCH page provides TE sequences for 315 species (Figure 5A), the CLASSIFICATION page allows online TE classification based on user-inputted sequences (Figure 5B), and the DOWNLOAD page offers ways to download reports, TEs, and pre-trained models (Figure 5C). The specific descriptions of these three modules are as follows:

**Figure 5.**
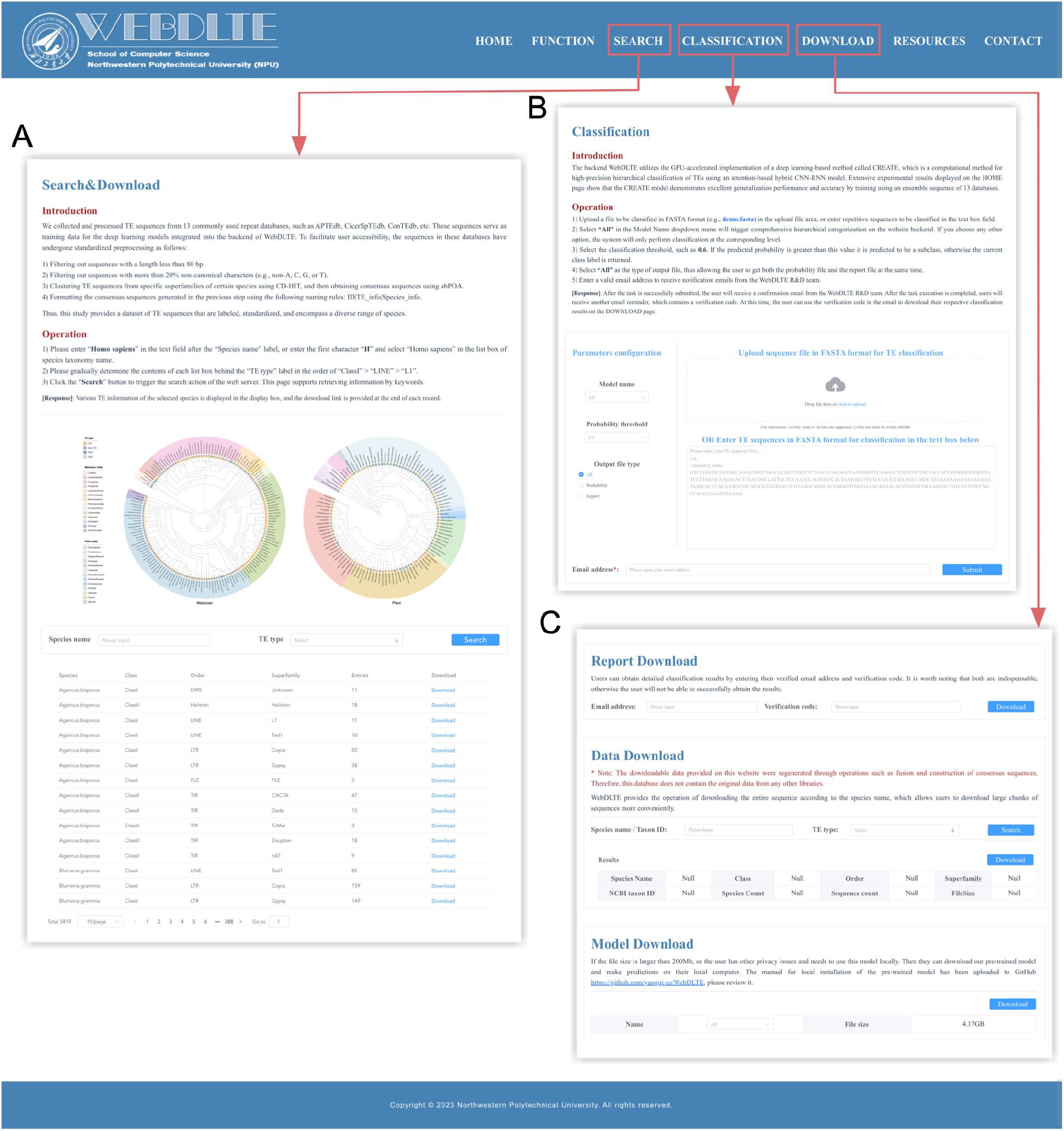
The main functions in WebDLTE. (A) Search function provides users to retrieve and download re-fused and re-processed TE consensus sequences for diverse species. (B) Users can upload repetitive sequences in FASTA format and provide their personal email address on the classification page. (C) The download page offers three functions, report download, data download and model download.

i. **Search and download TEs from 315 species**. WebDLTE is currently the most extensive TE database for eukaryotes, encompassing conserved TE sequences from 315 species, including animals, plants, and fungi. On the SEARCH page, users can retrieve and download re-fused and re-processed TE consensus sequences for all 315 species (Figure 5A). These sequences serve as valuable resources for conducting in-depth research on TEs.
ii. **Online classification of TE sequences**. WebDLTE offers an online service for TE classification, enabling users to conveniently upload repetitive sequences in FASTA format on the CLASSIFICATION page and provide their personal email address (Figure 5B). After the file is successfully submitted to WebDLTE’s backend, users will receive an email from the WebDLTE R&D team. Once WebDLTE completes processing of the submitted file, an email reminder will be sent again with a verification code. User can retrieve the classification results through the verification code in this email.
iii. **Download personal TE classification reports, pre-trained models, or TE consensus sequences**. The DOWNLOAD page provides three functions (Figure 5C). The Report Download module allows users to download the TE classification results by entering their email address and verification code. For users who need to perform arge-scale TE classification on their local systems, the pre-trained models can be downloaded via the Model Download module. Additionally, the Data Download module provides a variety of options for downloading re-fused and re-processed TE consensus sequences.

## 4. Discussion

Unlike other TE classification methods, deep learning is a concise and universal technique, which do not require prior experience or predefined features. CNN is a prominent deep learning model owing to its excellent feature extraction capabilities on high-dimensional data, and it is appropriate for revealing underlying sequence patterns. Meanwhile, RNN is another popular model that has shown efficient performance for sequential data. For deep learning-based approaches, feature extraction is an essential step to convert sequences into equal-length input vectors. *K-mer* counting is a commonly used method for feature representation, and it was also employed as a strategy to capture features from input sequences in our model. From another perspective, the flanking region of transposons contains a great variety of information about the properties and characteristics of TE types. Therefore, the *k-mer* counts were used as input vectors of the CNN module, and the handcrafted flanking sequences were fed as input features of the RNN module. Furthermore, an attention mechanism was employed to incorporate the features produced by CNN and RNN modules.

For each model in the class hierarchy, CTEATE outperformed other methods in different species. Separate experiments further demonstrated the effectiveness of the attention mechanism used in CTEATE. One interesting finding is that CTEATE-RNN presented excellent classification capacity in several TE classes, such as Class II sub1 and TIR models. This can attribute to the advantages of the handcrafted flanking sequences, which can represent structural motifs underlying TEs. Another intriguing finding is that CREATE-RNN also performed better in some minority classes than other CNN-based tools, which further enhanced the performance of CTEATE, e.g., TcMar and hAT in the TIR model, SINE1/7SL and SINE3/5S in the SINE model. Although these results are promising, additional efforts are required to address imbalanced data for enhanced outcomes.

For hierarchical classification, CTEATE got the best performance in different metrics compared with other methods. These experiments implied the probability threshold has a limited impact on the performance of CTEATE. It is also recommended to set this parameter to a higher value, such as 0.90, in practical applications. Furthermore, the results of this study may help us to investigate structural features in different TEs and the phylogenetic relationship between different transposition events. Nevertheless, this study trained each classifier separately, and the high-level classifier does categorically exclude some classes in lower levels.

WebDLTE provides the benefit of performing high-precision online categorization of TE sequences, utilizing the pre-trained model created by the CREATE method. This platform is equipped with query and download functions, enabling users to effortlessly retrieve and download sequences of interest. Without any limitations, users can perform searches by species name or transposon type. This innovative platform simplifies sequence annotation for researchers, opening new possibilities for investigating phylogenetic relationships. Moreover, WebDLTE serves as an extensive database for regenerated TE sequences, allowing users to easily access and download their preferred sequences. In addition, the platform offers a feature for analyzing the evolutionary relationships of transposons, which is beneficial for studying biological evolution.

## Data availability

The source code and demo data of CREATE are available at https://github.com/yangqi-cs/CREATE. The web interface of WebDLTE can be accessed at https://www.webdlte.nwpu.edu.cn. This website is freely accessible to all users without requiring any login or password.

## Author contributions

X.Q.S. supervised the project. Y.Q., X.Q.S., and X.Y.L. conceived and designed this work. Y.Q., X.Y.L., and Y.Q.C. conceived, design, and implemented the CREATE pipeline. X.Y.L., Y.Q., Y.F.W., Y.Y.L., and Y.Q.C. designed, and implemented the frontend and backend of WebDLTE. Y.Q., and Y.Q.C. preprocessed the data and coordinated data release. Y.Q., X.Y.L., M.H.G., and F.H.Z. analyzed the performance of algorithms developed in this study. Y.Q., and X.Y.L. wrote the paper. All authors have read and approved the final version of this paper.

